## Supplementary material for "Positive selection within the genomes of SARS-CoV-2 and other Coronaviruses independent of impact on protein function": Table 1

**Table 1. Coronavirus Accessions.**

| **Coronavirus Species** | **Name Used** | **NCBI Reference Sequence** | **Link** |
| --- | --- | --- | --- |
| Severe acute respiratory syndrome coronavirus 2 isolate Wuhan-Hu-1, complete genome | SARS-CoV-2 | NC_045512.2 | <https://www.ncbi.nlm.nih.gov/nuccore/NC_045512> |
| Bat coronavirus RaTG13, complete genome | Bat-CoV-RaTG13 | MN996532.1 | <https://www.ncbi.nlm.nih.gov/nuccore/MN996532.1> |
| Pangolin coronavirus isolate MP789, complete genome | Pan-CoV-GD | MT121216.1 | <https://www.ncbi.nlm.nih.gov/nuccore/MT121216.1> |
| Pangolin coronavirus isolate PCoV_GX-P4L, complete genome | Pan-CoV-GX | MT040333.1 | <https://www.ncbi.nlm.nih.gov/nuccore/MT040333.1> |
| Rhinolophus affinis coronavirus isolate LYRa11, complete genome | Bat-CoV-LYRa11 | KF569996.1 | <https://www.ncbi.nlm.nih.gov/nuccore/KF569996.1> |
| SARS coronavirus Tor2, complete genome  NCBI Reference | SARS-CoV | NC_004718.3 | <https://www.ncbi.nlm.nih.gov/nuccore/30271926> |
| Bat coronavirus BM48-31/BGR/2008, complete genome  NCBI Reference | Bat-CoV-BM48 | NC_014470.1 | <https://www.ncbi.nlm.nih.gov/nuccore/NC_014470.1> |
