## Supplementary info, other tables and figures for "Positive selection within the genomes of SARS-CoV-2 and other Coronaviruses independent of impact on protein function"

| | SARS-CoV-2 vs RATG13 | | | SARS_CoV-2 vs SARS-CoV | | | RATG13 vs SARS-CoV | | | Ancestral Seq vs SARS-CoV-2 | | | Ancestral Seq vs RATG13 | | | Ancestral Seq vs SARS-CoV | | |
|---|---|---|---|---|---|---|---|---|---|---|---|---|---|---|---|---|---|---|
| | RMSD (Å) | P-value | No. Twists | RMSD (Å) | P-value | No. Twists | RMSD (Å) | P-value | No. Twists | RMSD (Å) | P-value | No. Twists | RMSD (Å) | P-value | No. Twists | RMSD (Å) | P-value | No. Twists |
| nsp4 (whole sequence) | 15.77 | 1.43E-02 | 5 | 6.15 | 2.52E-03 | 5 | 12.64 | 2.94E-02 | 5 | 15.84 | 2.26E-02 | 4 | 19.34 | 4.96E-02 | 5 | 12.26 | 1.87E-02 | 5 |
| nsp4 (final 100 AA) | 1.53 | 1.11E-16 | 0 | 2.12 | 1.61E-13 | 0 | 1.9 | 3.89E-15 | 0 | 1.82 | 4.44E-16 | 0 | 1.96 | 1.11E-15 | 0 | 2.13 | 3.65E-14 | 0 |
| nsp5-3CL-pro | 0.6 | 0.00E+00 | 0 | 0.22 | 0.00E+00 | 0 | 0.29 | 0.00E+00 | 0 | 0.33 | 0.00E+00 | 0 | 0.34 | 0.00E+00 | 0 | 0.3 | 0.00E+00 | 0 |
| nsp16 | 0.17 | 0.00E+00 | 0 | 0.31 | 0.00E+00 | 0 | 0.33 | 0.00E+00 | 0 | 0.19 | 0.00E+00 | 0 | 0.17 | 0.00E+00 | 0 | 0.86 | 0.00E+00 | 0 |

**Supplementary Table 1.** Protein structures of Nsp4, Nsp5, and Nsp16 were predicted by PHYRE2. Pairwise comparisons of structure similarity by FATCAT shows all were significantly similar.

| | NC045512.2 vs MT517432 | | |
|---|---|---|---|
| | RMSD (Å) | P-value | No. Twists |
| nsp12 | 0.53 | 0.00E+00 | 0 |

**Supplementary Table 2.** Nsp12/RdRp protein structures for the Wuhan reference sequence (NC045512.2) compared to a sequence with the P323L mutation (MT517432) are significantly similar. Protein structures were predicted by PHYRE2 and compared using FATCAT.

| **Species** | **Strand (Forward or Reverse Complement)** | **minimum free energy prediction (kcal/mol)** | **free energy of the thermodynamic ensemble (kcal/mol)** | **ensemble diversity** |
|---|---|---|---|---|
| SARS-CoV-2 | F | -245.80 | -261.94 | 201.42 |
| | RC | -204.00 | -222.12 | 214.17 |
| RaTG13 | F | -246.40 | -263.02 | 195.65 |
| | RC | -208.30 | -227.52 | 156.88 |
| SARS-CoV-2/RaTG13 Ancestral Reconstruction | F | -246.40 | -261.86 | 189.39 |
| | RC | -210.60 | -227.35 | 160.77 |
| Pan-CoV-GD | F | -231.70 | -247.34 | 246.94 |
| | RC | -208.20 | -224.21 | 182.69 |
| SARS-CoV | F | -255.50 | -271.43 | 189.59 |
| | RC | -241.00 | -256.39 | 161.73 |

**Supplementary Table 3.** NSP16 RNA free energy predictions and ensemble diversity calculated by RNAFold Web Server.

### Supplementary Figures

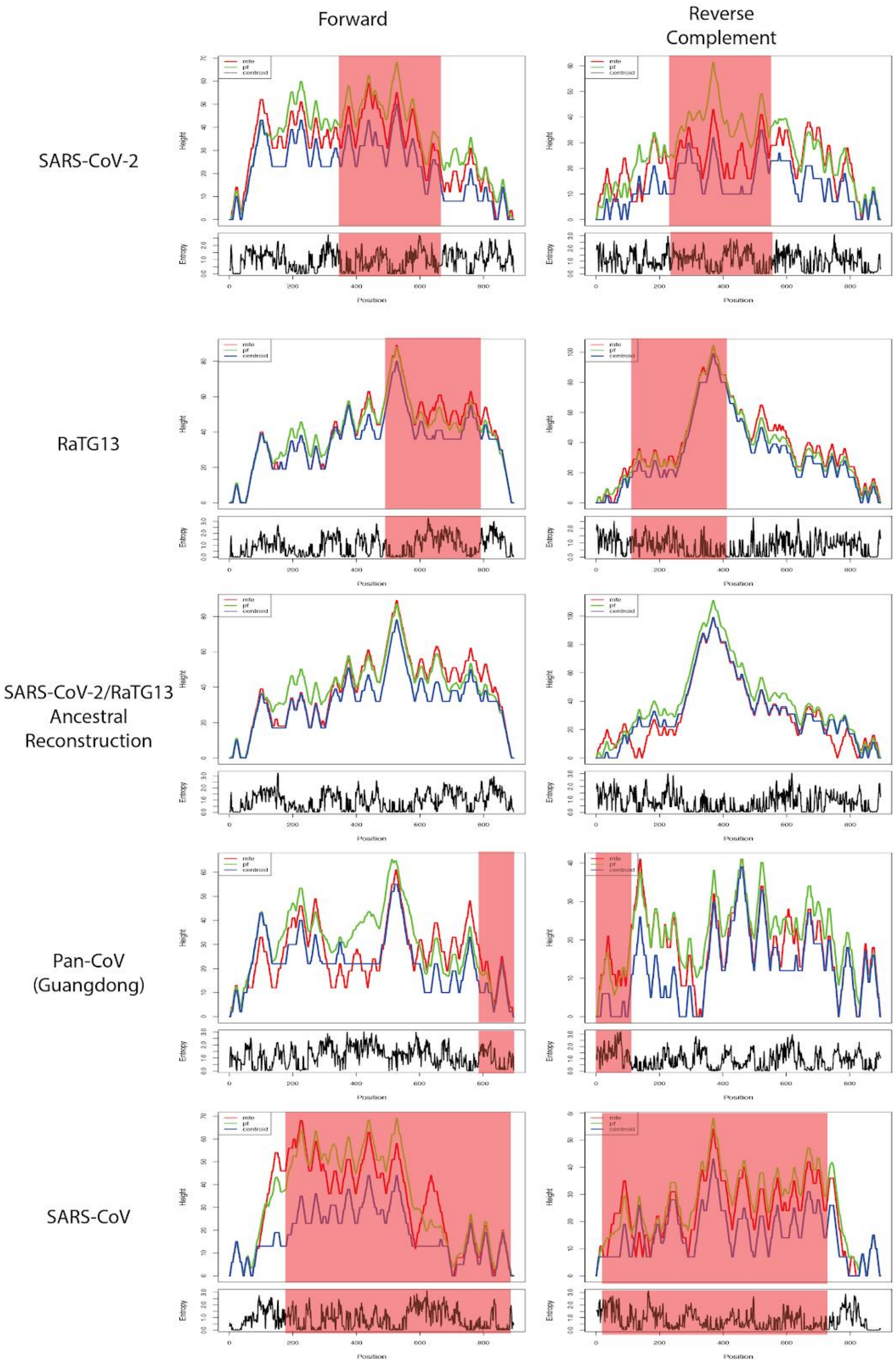


**Supplementary Figure 1.** MFE mountain plots for the forward and reverse strands of Nsp16 in SARS-CoV-2, Bat-CoV-RaTG13, the reconstructed ancestor of SARS-CoV-2 and RaTG13, Pan-CoV-GD and SARS-CoV at 37 °C. Regions under positive selection shaded in red.

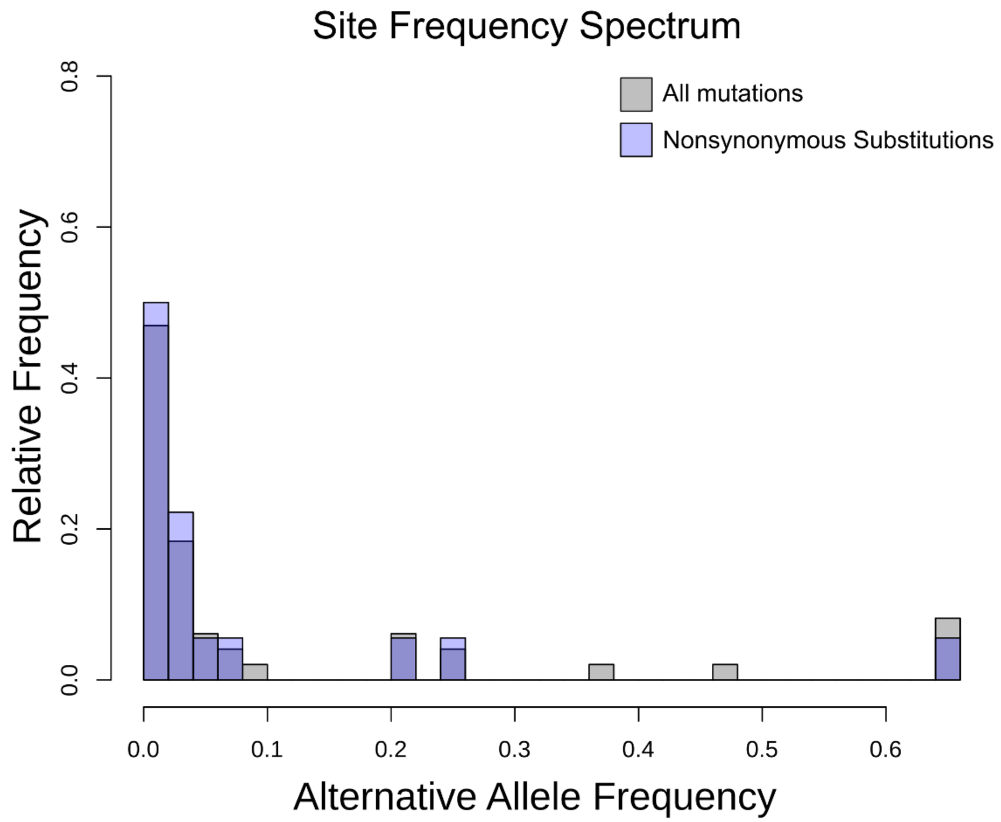


**Supplementary Figure 2. Site Frequency Spectrum.**

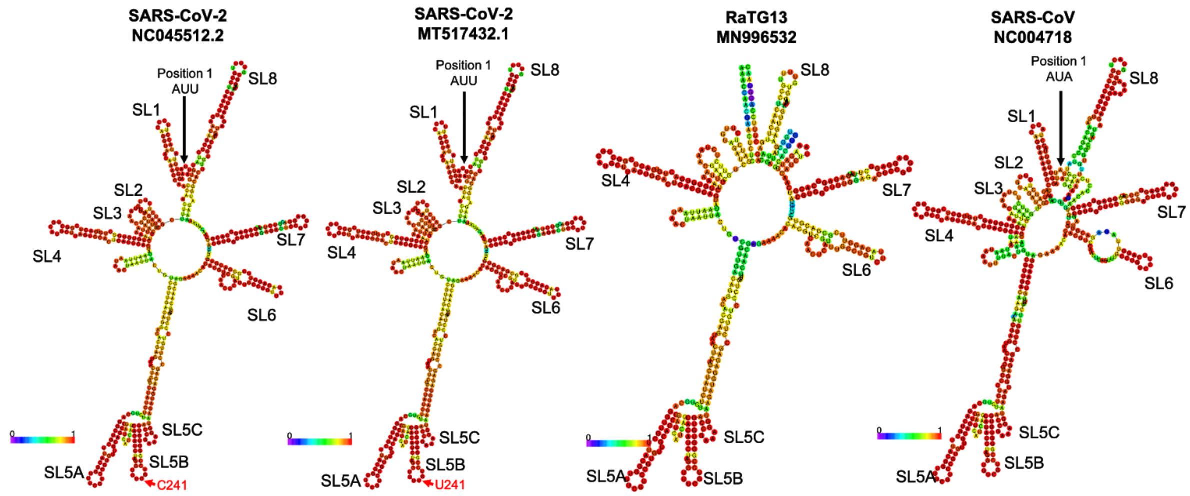


**Supplementary Figure 3.** Secondary structure of 5' UTR (first 474 bp) predicted by RNAfold. The C>U mutation at position 241 is within the SL5B stem-loop structure, indicated with a red arrow. The Pan-CoV-GD 5' UTR sequence is missing the first 129 bp relative to the Wuhan SARS-CoV-2 reference sequence and so is not included here.
